# Stress-mediated convergence of splicing landscapes in male and female Rock Doves

**DOI:** 10.1101/798413

**Authors:** Andrew S Lang, Suzanne H Austin, Rayna M Harris, Rebecca M Calisi, Matthew D MacManes

## Abstract

**Background:** The process of alternative splicing provides a unique mechanism by which eukaryotes are able to produce numerous protein products from the same gene. Heightened variability in the proteome has been thought to potentiate increased behavioral complexity and response flexibility to environmental stimuli, thus contributing to more refined traits on which natural and sexual selection can act. While it has been long known that various forms of environmental stress can negatively affect sexual behavior and reproduction, we know little of how stress can affect the alternative splicing associated with these events, and less still about how splicing may differ between sexes. Using the model of the rock dove (*Columba livia*), our team previously uncovered sexual dimorphism in the basal and stress-responsive gene transcription of a biological system necessary for facilitating sexual behavior and reproduction, the hypothlamic-pituitary-gonadal (HPG) axis. In this study, we delve further into understanding the mechanistic underpinnings of how changes in the environment can affect reproduction by testing the alternative splicing response of the HPG axis to an external stressor in both sexes.

**Results:** This study reveals dramatic baseline differences in HPG alternative splicing between males and females. However, post submitting subjects to a restraint stress paradigm, we found a significant reduction in these differences between the sexes. In both stress and control treatments, we identified a higher incidence of splicing activity in the pituitary in both sexes as compared to other tissues. Of these splicing events, the core exon event is the most abundant form of splicing and more frequently occurs in the coding regions of the gene. Overall, we observed less splicing activity in the 3’UTR end of transcripts than the 5’UTR or coding regions.

**Conclusions:** Our results provide vital new insight into sex-specific aspects of the stress response on the HPG axis at an unprecedented proximate level. Males and females uniquely respond to stress, yet exhibit splicing patterns suggesting a convergent, optimal splicing landscape for stress response. This information has the potential to inform evolutionary theory as well as the development of highly-specific drug targets for stress-induced reproductive dysfunction.

## Introduction

Organismal behavior and its mechanistic underpinnings have been consistent quandaries for many biologists (Vermeij, 1991; Sutherland, 1998; Holway & Suarez, 1999; Wong & Candolin, 2015). Even among the few cases in which an adaptive behavior is clearly understood, we have little information on how proximate mechanisms have led to ultimate behavioral adaptations. One of the mechanisms by which animals modulate their response to stimuli is by adjusting the levels of endogenous protein products in physiological systems. (Sapolsky, Romero & Munck, 2000; Zhao et al., 2017). This variable production may result in differential binding of hormones and signaling peptides, enabling an organism to receive information more accurately about external stimuli and effectively react (Ohama et al., 2017; Calisi et al., 2018). In addition to adjusting the quantity of a given protein, organisms may further alter these proteins by producing slightly modified versions (e.g., isoforms) of each transcript (Smith, Patton & Nadal-Ginard, 1989; Tapial et al., 2017). This variable modification of gene products is called alternative splicing.

Alternative splicing is a process unique to eukaryotes (Izquierdo & Valcárcel, 2006) that involves cleavage of transcribed RNA at specific splice sites and varying inclusion or exclusion of genomic elements (introns and exons). In the human genome, approximately 80% of exons are >200bp in length (Krawczak et al., 2007; Gudlaugsdottir et al., 2007); however, exon sizes identified in other species vary from a single base to >17,000bp in length. Each human gene contains, on average, eight exons (Sakharkar, Chow & Kangueane, 2004). This variable inclusion of genetic sequences results in a dramatic increase in the number of potential transcript and protein products that a single gene may produce. Alternative splicing presents an additional mechanism by which mRNA levels and gene expression can be regulated, while also greatly increasing proteome diversity. Splicing activity is thought to be responsible for the majority of proteomic diversity in eukaryotes (Birzele, Csaba & Zimmer, 2008) and, potentially, may be an underlying mechanism of functional genomic evolution (Bush et al., 2017).

Numerous types of splicing events exist that occur at different frequencies in a given genome and alter proteins in subtle to dramatically different ways (Kim, Magen & Ast, 2007). Cassette exon splicing (also referred to as exon skipping) is the most common type of splicing event in vertebrates and invertebrates, while intron retention is more common in plants. Additionally, alternative selection of 5’ and 3’ splice sites, coupled with variable adenylation of the transcript, results in further modification of protein products (Gueroussov et al., 2015; Tian & Manley, 2017). The splicing process consists of two major steps: assembly of the spliceosome and the actual splicing of pre-mRNA (Wang et al., 2015). In brief, the spliceosome is comprised of several small nuclear ribonucleoproteins that positionally establish the 5’ splice site, the branch point sequence, and the 3’ site. An assembly of spliceosome complexes and eight evolutionarily-conserved RNA-dependent ATPases/helicases is then followed by the execution of numerous splicing steps, ultimately resulting in exon excision, exon ligation, or intron retention (Wang et al., 2015). The inclusion of an exon in the final mRNA product is entirely driven by cis- and trans-acting elements/factors. The interaction of these elements within the splicing process promotes or inhibits spliceosome activity on various splice regions, resulting in alternative splicing (Wang et al., 2006; Wang & Burge, 2008).

Alternative splicing mechanisms enable organisms to sense and react to minute changes in the local environment, allowing both plants and animals to tailor their responses to their surroundings with extreme precision (Staiger & Brown, 2013; Preußner et al., 2017). Previous research has revealed unique roles for alternative splicing in the immune response of chickens with avian pathogenic E.coli (Sun, 2017), mediation of abiotic stress response pathways of plants (Laloum, Martín & Duque, 2018), and enhanced fear memory of mice (Nijholt et al., 2004). Alternative splicing has also been implicated in various aspects of cancer, including oncogenesis (Climente-Gonzalez et al., 2017) and cancer drug resistance (Chen & Weiss, 2015; Siegfried & Karni, 2018). Some studies have identified a sex-bias in alternative splicing in Drosophila (McIntyre et al., 2006; Telonis-Scott et al., 2009; Gibilisco et al., 2016), while others have identified unique sex-specific splicing differences in human brains (Trabzuni et al., 2013). The diverse roles of alternative splicing in biological processes and behavioral responses inherently speak to the depth and breadth that alternative splicing drives organismal physiology and behavior, at both local and global levels. By identifying the splicing landscape that modulates gene expression and mRNA transcript composition in both males and females, we increase the resolution at which we can comprehend the proximate mechanisms underlying animal physiology and behavior.

In vertebrates, a symphony of physiological events is required to regulate sexual behavior and reproduction, and these mechanisms are driven by an interconnected biological system made up of the hypothalamus in the brain, the pituitary gland, and the gonads (testes/ovaries) (Sower, Freamat & Kavanaugh, 2009; Nozaki, 2013; MacManes et al., 2017; Calisi et al., 2018). This hypothalamic-pituitary-gonadal (HPG) axis can be disrupted in multiple, complex ways (Johnson et al., 1992; Toufexis et al., 2014; Geraghty & Kaufer, 2015; Calisi et al., 2018). However, we know little about how stress affects the HPG axis at the level of alternative splicing, and we know even less regarding its effects at this level in males versus females. Understanding how the alternative splicing landscape of the reproductive axis changes in the face of stress will not only offer more insight into how stress can affect reproduction, but deepen our proximate knowledge of biological processes and sexually-biased behavioral responses in general.

Using the classic reproductive model (Darwin, 1860; Lehrman, 1955; Ball & Silver, 1983; Buntin, Ruzycki & Witebsky, 1993) and rising genomics model (Shapiro et al., 2013; Gillespie et al., 2013; Domyan et al., 2016; MacManes et al., 2017; Calisi et al., 2018) of the rock dove, *Columba livia*, we have identified sexual dimorphism in both basal (MacManes et al., 2017) and restraint stress-responsive (Calisi et al., 2018) HPG gene expression at the level of RNA transcription. In this study, we traverse beyond the level of transcription to test for sex-biased alternative splicing patterns in the HPG axis of the rock dove in response to a restraint stress stimulus. Using a relatively highly-replicated (n=12/sex) study design, we identify significantly similar and different splicing events between the sexes and in response to restraint stress treatment. To our knowledge, this is the first report of sex-specific splicing events in the HPG axis in response to a stressor.

## Materials and Methods

### Animal Collection

We housed birds at the University of California, Davis, in semi-enclosed aviaries (5’ x 4’ x 7’), with 8 sexually reproductive adult pairs per aviary. Food and water were provided *ad libitum*. Birds were exposed to natural light, which was supplemented with artificial light on a 14L:10D cycle. We sampled sexually mature males and females that were without eggs or chicks so as to control for physiological changes that occur to facilitate parental care behaviors. Birds were sampled between 0900—1100 (PST) to also control for potential circadian rhythm confounds. All handling and sampling procedures followed approved animal care and handling protocols (UC Davis IACUC 18895). Samples from the control group were taken within 5min of entering their cages. Birds in the restraint stress treatment group were captured in ¡1min upon entering their aviary and immediately and individually restrained in cloth bags for 30min prior to sampling, replicating methods described in Calisi et al. 2018.

Prior to sampling, birds were anesthetized using isoflurane, at which point they were immediately decapitated. Trunk blood was collected for the detection of concentrations of the adrenal hormone, corticosterone; high concentrations confirmed the effectiveness of the stress treatment protocol (Calisi et al., 2018). All tissues (brains, pituitaries, and gonads) were flash frozen on dry ice and transferred to a - 80 °C freezer for storage until additional processing, a process described in (Shapiro et al., 2013; Gillespie et al., 2013; MacManes et al., 2017; Calisi et al., 2018). Briefly, a cryostat (Leica CM 1860) was used to section the brains coronally at 100 microns to enable optimal visualization and biopsy of the hypothalamus, which included the adjoined lateral septum. Hypothalamic sections, pituitaries, and gonads were preserved in RNALater (Invitrogen, Thermo Fisher Scientific) and shipped on dry ice from UC Davis to the University of New Hampshire for cDNA library preparation and sequencing. We sequenced tissue from whole homogenized hypothalami, pituitaries, testes and (homogenized tissue from the oviduct and ovarian follicles). In total, we processed and sequenced hypothalami, pituitary glands, and gonads (testes/ovaries) from 48 birds (12 males and 12 females, each, per treatment —control and stress), resulting in 144 cDNA libraries.

### cDNA Library Preparation and Sequencing

Library preparation and sequencing has been previously described in (Shapiro et al., 2013; Gillespie et al., 2013; MacManes et al., 2017; Calisi et al., 2018). In brief, all tissue samples were thawed on ice in an RNAse-free work environment. Total RNA was extracted with a standard Trizol extraction protocol (Thermo Fisher Scientific, Waltham, MA). Total RNA quality and quantity was assessed with gel electrophoresis and a Broad Range RNA Qubit assay (Thermo Fisher Scientific, Waltham, MA). Illumina sequence libraries were prepared using the NEB Next Ultra Directional RNA Library Prep Kit (New England Biolabs, Ipswich, MA). Quality and quantity of the cDNA libraries were validated with a Tapestation 2200 Instrument (Agilent, Santa Clara, CA). Libraries were diluted to 5nM and pooled. We sent multiplexed library pools to the New York Genome Center for 125 base pair paired-end sequencing on a HiSeq 2500 platform. 16 samples were re-sequenced at Novogene Sequencing Center to increase coverage of those samples with the lowest read abundance.

### Pigeon Genome Annotation

Many of the currently-available methods for identification of splicing events are heavily dependent upon genome assembly completeness and annotation quality. The existing annotated assembly for *C. livia* (GCF 000337935.1) did not provide the level of contiguity required for accurate identification of splicing events. To address this, we used an unannotated chromosome-level assembly for the rock dove (GCA 001887795.1). We annotated this chromosome-level assembly with Maker version 2.3.1(Cantarel et al., 2008). A custom script (available here: https://github.com/AndrewLangvt/Dissertation/blob/master/cliv2_gff2gtf.py was used to generate the required GTF file from the resultant GFF.

### Sequence QC, Read Processing, and Splicing Event Identification

Sequence data were downloaded to the Pittsburg Supercomputing Center network “Bridges” and read quality confirmed with FastQC v0.11.5 (Andrews, 2011). Read data from all 144 samples was first error corrected with Rcorrector v3 (Song & Florea, 2015), then adapter sequences and corrected reads with PHRED <2 were removed from sequence datasets.

To accurately identify splicing events in our study system, we used Whippet, an algorithm that rapidly identifies splicing events from RNAseq data (Sterne-Weiler et al., 2017). (Code available here: https://github.com/AndrewLangvt/Scripts/tree/master/splicing_analysis/whippet). Whippet uses annotated transcript features to generate splice graphs and transcript-level mapping data to calculate Percent Spliced In (PSI) values for all paths through the directed graph (Sterne-Weiler et al., 2017). Whippet v0.10.4 was used to quantify read data and identify differential splicing events between treatments. Events with probability >0.95 and abs(deltaPSI) great than or equal to 0.1 were deemed significant and considered to be true splicing events.

### Splicing Event Comparisons

To understand the sex-specific response to restraint stress, we asked (1) how similar are males and females at the exonic level (i.e. assessing splicing events between sexes, within treatments), and (2) how do males and females differ in their response to stress (i.e., assessing splicing events within the sexes, between treatments). We did not include analyses between different tissues (e.g., male gonads to female gonads, or male pituitary to male hypothalamus) as numerous other studies have identified tissue-specific splicing events; these comparisons would not aid in discerning unique splicing patterns between sexes, or between treatments (Grosso et al., 2008; Saha et al., 2017; Pineda & Bradley, 2018). Whippet identifies splicing events by type, including Core Exon (CE), Alternate Acceptor (AA), Alternate Donor (AD), Retained Intron (RI), Tandem Alternative Polyadenylation Site (TE), Tandem Transcription Start Site (TS), Alternative First Exon (AF), and Alternative Last Exon (AL). Of all the event types detected by Whippet, core exon, alternate acceptor, alternate donor, and retained intron encompassed 95% of our identified splicing events. For simplicity, we did not represent the TE, TS, AF, or AL events in our figure depicting splicing by type even though these events are included in all of the analyses.

### Gene Ontology Parent Term Analysis

We assessed abstracted (or parent) gene ontology (GO) terms of spliced genes to identify patterns of enrichment or depletion of gene functions. Lists of Entrez Gene IDs were analyzed through the “Gene List Analysis” tool at http://www.pantherdb.org to link GO terms with each ID. These terms were further analyzed with go abstract.py (https://github.com/twestbrookunh), a python script that abstracts a provided ontology term to its “parent” term from the Gene Ontology Consortium Database (Ashburner et al., 2000; Mi et al., 2017; The Gene Ontology Consortium, 2019). We generated a matrix of expected abundances for each tissue using this same analysis on the entire C. livia genome. GO terms with counts deemed significantly different from expected counts via Chi-Squared analyses were visualized in R. We calculated differences of counts from expected values and did not include parent ontology terms of very low abundance in the genome (<2%) in our visualization.

### Exon Size, Location, and Motif Analysis

We used a custom Python script, available at https://github.com/AndrewLangvt/Scripts/blob/master/splicing_analysis/exon_extract.py to extract desired exons from the genome and translate them into amino acid sequences based upon the phase and strand delineated in the GFF while simultaneously determining the length of each exon and the transcript region from which it was spliced (5’-UTR, CDS, or 3’-UTR). Distributions of exon lengths were compared to the genomic distribution to determine statistical significance using a Wilcoxon test. Expected counts of splicing events per region were determined from the genomic proportions and difference from expected was normalized by expected values across tissues. We further assessed the functional nature of alternatively spliced exons by investigating these sequences for protein motifs. To accomplish this, we used MOTIF, a GenomeNet tool (https://www.genome.jp/tools/motif/) which identifies protein motifs within protein sequences by searching the PFAM database (Finn et al., 2016) and NCBI databases COG (PMID: 10592175), SMART (PMID: 10592234), and TIGRFAM (PMID: 11125044).

## Results

### Sequencing Results, Read Data, and Code Availability

Samples were sequenced to a read depth between 2.3 million and 24.5 million read pairs, for a total of 144,363,425 paired-end reads (more fully described in (Calisi et al., 2018). Read data corresponding to the control birds are available using the European Nucleotide Archive project ID PRJEB16136; read data corresponding to the stressed birds are available at PRJEB21082. All code for analyses in this manuscript can be found at https://github.com/AndrewLangvt/Scripts/tree/master/splicing_analysis/.

### Male vs. Female Splicing Comparison

Our first aim was to understand sex-typical splicing in the hypothalamus and pituitary by assessing each tissue for alternative splicing events between males and females. We counted the number, and type, of alternatively spliced loci between males and females in each treatment state (control: male vs female; stress: male vs female). This approach allowed us to determine how the splicing landscape changed between sexes in response to restraint stress, and in which state the sexes shared a more similar splicing profile. As previously stated, we did not include gonads in this comparison due to inherent splicing differences between tissue types. Chi-squared tests (hereafter, ChiSq) were used to determine statistical significance (p <0.05) throughout our analyses; all p-values, degrees of freedom, and sample sizes are included in Table 1.

**Table 1.**
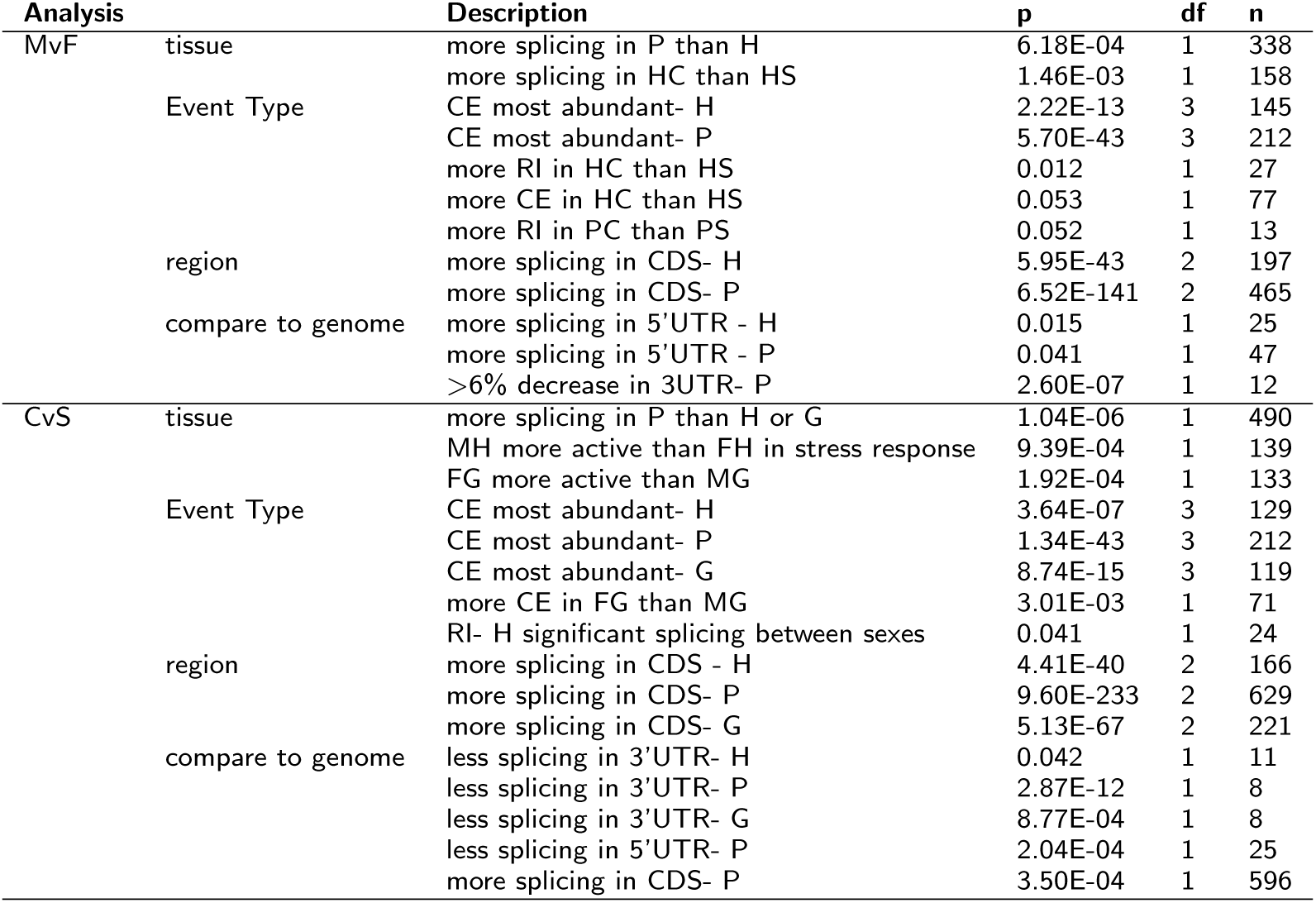
Statistics for all Chi-Square Tests. This table contains all Chi-Square values, degrees of freedom (df), and sample size (n) for every test of significant splicing events in this paper. H=Hypothalamus. P=Pituitary. G=Gonad. C=Control. S=Stress. CE=Core Exon. RI=Retained Intron. CDS=Coding Sequence. UTR=Untranslated Region.

### Male vs. Female Splicing Comparison: Events by Type

In total, we identified 158 splicing events in the hypothalamus and 225 events in the pituitary. When compared to the hypothalamus, the 42% increase of splicing event abundance seen in the pituitary is significant (ChiSq p=6.18e-4). In both tissues, more events were identified in the control state compared to the stress condition (hyp: 99 control/59 stress, pit: 123 control/102 stress), but only the relationship in the hypothalamus was statistically significant (Figure 1, ChiSq p: hyp=1.46e-3; pit=0.162).

**Figure 1.**
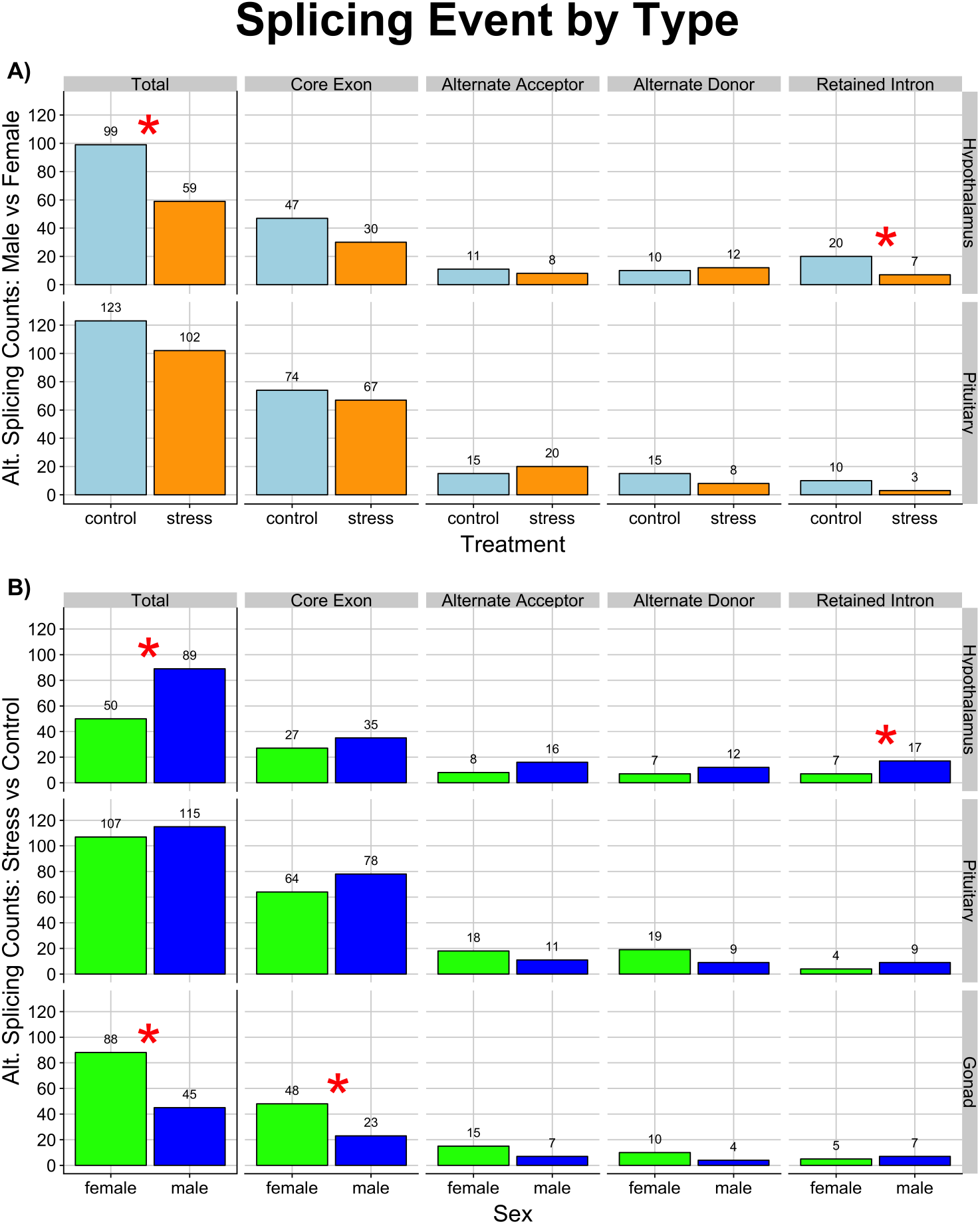
Splicing events by type for both the a) Male vs Female and b) Control vs Stress comparisons. Rows denote tissue type (labeled on the right), and counts of splicing events are further broken down by event type. Alternatively spliced genes in the male vs female analysis revealed, in both tissues, more events in the control versus restraint stress condition. The core exon event was the most abundant regardless of tissue or treatment. Light blue represents the control group; yellow is restraint stress. Hypothalamic retained intron events were the only event to differ significantly between treatments, represented by a red star (ChiSq p=0.012). In the control vs splicing comparison, more splicing occurred in the male hypothalamus; while in the gonad, more splicing occurred in the female. Blue represents males; green represents females. Red stars represent statistical significance between abundances in males and females, with more core exon splice events occurring in the female gonad than male (ChiSq p=3.01e-3), and more hypothalamic retained intron events found in males than females (ChiSq p=0.041).

These total counts were further broken down by event type (Figure 1). The core exon event was the most abundant event identified across sex in both the hypothalamus and pituitary, regardless of treatment (ChiSq p: hyp=2.22e-13, pit=5.70e-43). Of these core exon splice events, there were almost twice as many in the pituitary compared to hypothalamus (pit: 74 control/67 stress, hyp: 48 control/30 stress). Within these event types, we tested for statistical significance between splicing differences in males and females of each treatment group. Retained intron events in the hypothalamus were the only event to differ significantly between treatments (ChiSq p=0.012), with nearly 3 times (280% increase) more splicing events in the control state than the stressed. Both core exon events in the hypothalamus and retained intron events in the pituitary reflected a similar increased abundance in splicing events of the control state, though these relationships were not significant (ChiSq p: CE-Hyp=0.053, RI-Pit=0.052). The distribution of PSI values between males and females did not vary between treatments, indicating that the level of event inclusion/exclusion difference between the sexes was generally unaffected by treatment.

### Male vs. Female Splicing Comparison: Genes of Interest

Using our comparison of male to female splicing patterns, we were able to identify sex-specific alternatively spliced genes in the stress response. We provide a full list of spliced genes within each comparison (Table S1). Some of these spliced genes are involved in functional gene expression within the HPG axis. POU class 2 homeobox 1 (POU2F1), a transcription factor that regulates transcription of gonadotropin-releasing hormone (GnRH) (Chandran & DeFranco, 1999; Cheng et al., 2002), is alternatively spliced in the male pituitary stress response. GnRH is a primary regulator of the HPG axis (Pohl et al., 1983; Campbell, Dobson & Scaramuzzi, 1998; Takahashi et al., 2016). Splicing of POU2F1 likely affects the HPG axis, indirectly, by modulating transcription of GnRH (Rosen, Jameel & Barkan, 1988; Schneider, Tomek & Gründker, 2006; Jin & Yang, 2014). Through alternative splicing of the POU2F1 gene in the pituitary, males may be altering signaling pathways within the HPG axis to optimize stress response.

Few genes were consistently alternatively spliced between males and females in both treatments. Those that did exhibit consistent alternative splicing between the sexes were often related to immune function. Rap1 GTP-ase activating protein 1 (RAP1GAP) is consistently alternatively spliced between male and female hypothalami, in both the control and stress treatments. Previous findings have shown RAP1GAP to be a putative oncogene (Nellore et al., 2009). This gene mediates the strength of cell adhesions through regulation of Rap1, thus modulating T-cell response (Katagiri et al., 2002). Alternatively spliced between sexes in the pituitary, P-selectin (SELP) is known to preserve immune function in mice (González-Tajuelo et al., 2017). The corresponding ligand, P-selectin glycoprotein ligand-1 (PSGL-1), negatively regulates T-cell response through binding of SELP (Matsumoto, Miyasaka & Hirata, 2009). Genes consistently alternatively spliced between males and females may reflect splicing-level sexual dimorphism, indicating that males and females inherently differ in their splicing landscapes. Further, these differences appear to speak to a sex-specific stress response through modification of genes related to immune processes. Through future study of the splicing landscape of these genes of interest in additional tissues and states, we will likely reveal additional inherent splicing differences between the sexes.

### Male vs. Female Splicing Comparison: Gene Ontology

By observing abundances of parent ontology terms significantly deviating from genomic expectation, we were able to gain a broader understanding of gene-types targeted by alternative splicing. In the list of significant Molecular Function terms (Figure 2), splicing of organic and heterocyclic compound genes is underrepresented in the pituitary, while small molecular and drug binding is overrepresented in the hypothalamus regardless of treatment. There was an interaction seen in the hypothalamus; splicing of organic and heterocyclic compound genes was overrepresented in the stress treatment, though underrepresented in the control group. Biological Process terms suggest that males and females differ very little in their splicing profile of metabolic genes in either the hypothalamus or pituitary, given there were fewer spliced genes with metabolism GO terms in these tissues (Figure S1). Finally, splicing events in stressed males and females are more abundant in genes related to cell/neuronal structure of the pituitary than the hypothalamus (Figure S2).

**Figure 2.**
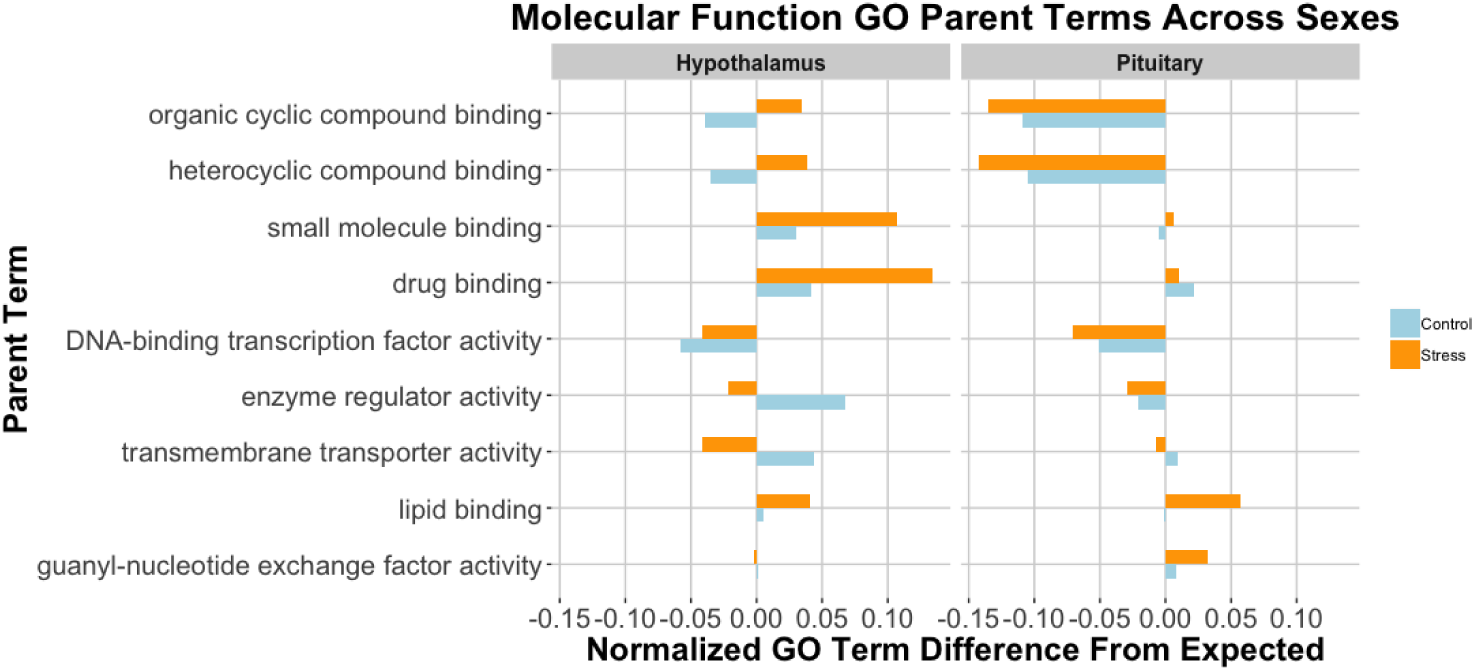
Molecular Function GO analysis, Male vs. Female (normalized counts of observed-expected). Splicing in the pituitary is more prevalent in heterocyclic and organic cyclic compound binding genes, while splicing in the hypothalamus affects small molecule and drug binding loci. Counts for all terms in this figure were significantly different from the expected value in at least one of the tissues. Parent ontology terms along the y-axis are in descending order from most frequent in the genome to less frequent. The left panel depicts counts from hypothalamic spliced genes, and the right panel spliced genes from the pituitary. Blue represents control treatment, and orange is restraint stress. Abundances are observed counts – expected (based upon genomic predictions)/ total events within that tissue. We did not include any terms that were attributed to less than 2% of the genome.

### Male vs. Female Splicing Comparison: Exon size, location, and motifs

To further characterize the splicing profile of the sexes, we visualized distributions of exon sizes, where these exons were located, and protein motifs contained therein. In terms of size distribution, our analyses revealed no significant difference between the control (male vs female) and stress (male vs female) comparisons in the hypothalamus or pituitary, suggesting that the size of alternatively spliced exons between males and females does not differ between control or stressed states (Figure 3). In all cases, spliced exons were smaller than predicted by the genomic distribution (Wilcoxon test p: hyp-control= 4.52e-12; hyp-stress= 2.16e-3; pit-control= 2.75e-8; pit-stress= 3.53e-10) (Figure 3). Splicing events occurred in a variety of protein motifs, but no particular motif was significantly more spliced than the others (Figure S3).

**Figure 3.**
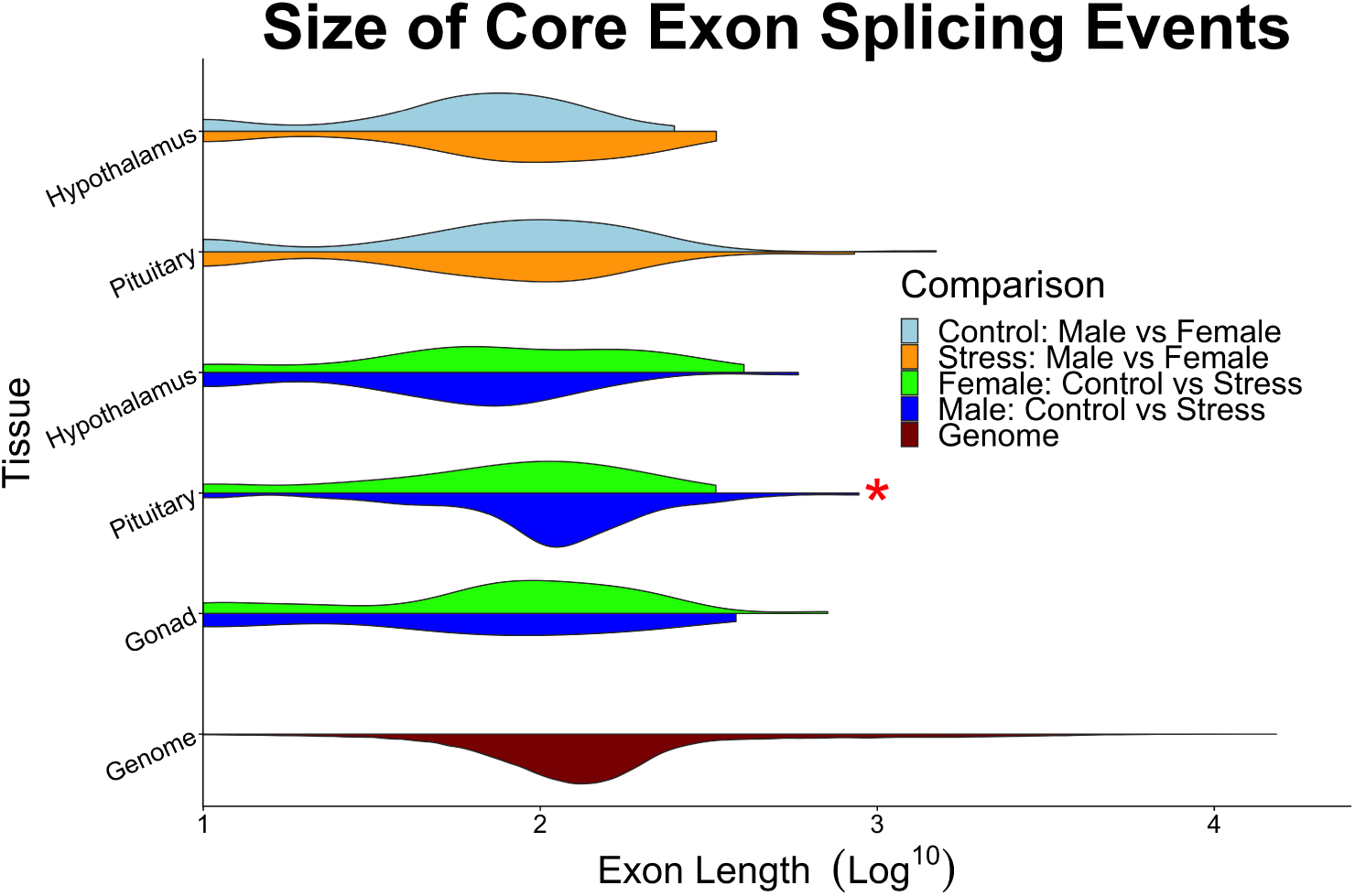
Distributions of core exon splicing event lengths for between-sex spliced loci in the control (light blue) and stress (orange) states as well as control-stress spliced loci in male (blue) and female (green) states. Neither the hypothalamus or pituitary showed significant difference between control (light blue) vs stress (orange) comparisons. In the pituitary of control vs stress comparison, lengths of spliced exons are significantly larger in the male pituitary than that of spliced exons in the female pituitary (Wilcoxon, p = 0.022). Distribution of genome exon sizes is colored in red. All comparisons with genome distribution, exon sizes were significantly smaller (Wilcoxon, p <2.80e-3).

In both tissues, core exon splice sites occurred in protein coding sequences much more frequently than either of the untranslated regions (ChiSq p: hyp=5.95e-43, pit=6.52e-141). This, perhaps, is not surprising given that the CDS regions are more abundant in the genome and alteration to these regions will ultimately result in changes to the protein sequence. The abundance of alternatively spliced exons present in the 5’ & 3’ untranslated regions of the hypothalamus and pituitary was significantly different from expected values (Figure 4). In the hypothalamus, more spliced exons occurred in the 5’UTR than genomic proportions would predict, with more dramatic shifts in the restraint stress treatment than the control (ChiSq, p=0.015). In the pituitary, the control group exhibited more spliced exons of the 5’UTR and both treatment groups presented more than a 6% decrease of splicing events in 3’UTR regions than predicted from genomic values (ChiSq p: 5’UTR=0.041, 3’UTR=2.60e-7).

**Figure 4.**
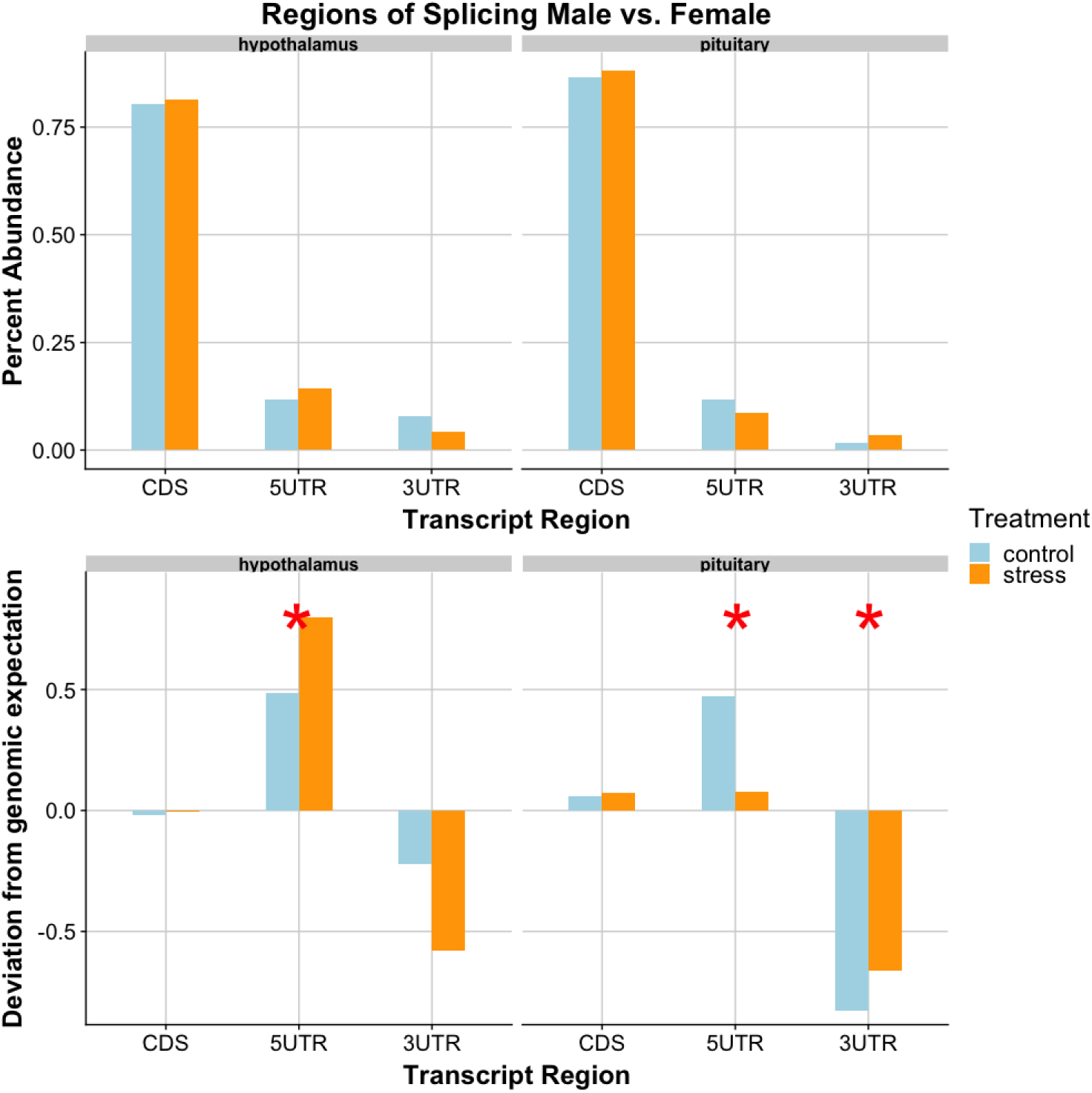
Transcriptional regions of male-female splicing events. In both tissues, the 5’UTR region was a site of increased splicing activity, while the 3’UTR regions underwent less splicing. The upper half represents percentage of total splicing events (for each tissue) in each transcript region. There were significantly more CDS events than 5UTR or 3UTR (ChiSq p: hyp=5.95e-43, pit=6.52e-141). The lower portion depicts how these counts differ from the expected (i.e. a value of 0.5 here indicates a 50% deviation from genomic expectation). Blue represents control treatment, while orange is restraint stress. Red stars represent significance between observed counts and expected from genomic predictions (ChiSq p: Hyp-5’UTR=0.015, Pit-5’UTR=0.041,Pit-3’UTR=2.60e-7).

### Control vs. Stress Splicing Comparison

The second aim of this study was to observe splicing differences within each sex in control and stress states, to identify how each sex individually responded to restraint stress. We compared, within each sex, control and stress treatments (female: control to stress; male: control to stress). We counted the number of alternative splicing events for these within-sex, across-treatment comparisons. Here, we include comparisons between gonads as the alternative splicing events identified are within-sex and thus, we can observe how male and female gonads individually respond to restraint stress. ChiSq tests were used to determine statistical significance throughout our analyses; all p-values, degrees of freedom, and sample sizes are included in Table 1.

### Control vs. Stress Splicing Comparison: Events by Type

Similar to our comparison of male-female alternative splicing, control-stress splicing reveals more splicing in the pituitary than other tissues (ChiSq p=1.04e-6). The male hypothalamus is more active in stress-response splicing compared to the female hypothalamus (ChiSq p=9.39e-4), and the female gonad exhibits more activity than the male gonad (ChiSq p=1.92e-4). The male hypothalamus displays 59% more splicing events than the female hypothalamus, and we identified 78% more splicing events in the ovaries than the testes (Figure 1).

Of all event types, the core exon event was most abundant in all tissues (ChiSq p: hyp=3.64e-7, pit=1.34e-43, gon=8.74e-15)(Figure 1). Of these core exon events, the gonads were the only tissue to present statistical significance across sexes; more core exon splice events occurred in the ovaries than in the testes (ChiSq p=3.01e-3). This parallels our previous findings of elevated female gonadal gene expression in response to restraint stress (Calisi et al., 2018). The only other event that differed significantly between the sexes was hypothalamic retained intron events, with more found in males than females (ChiSq p=0.041).

### Control vs. Stress Splicing Comparison: Genes of Interest

Through assessment of alternative splicing events between control and stress states, we were able to identify genes spliced in response to stress within each sex. Estradiol 17-beta-dehydrogenase 11 (HSD17B11) was alternatively spliced between treatments in the male hypothalamus. HSD17B11 plays a role in hormone metabolism, through which it may mediate endogenous estrogen levels (Ariazi et al., 2011). Through feedback on the brain, estrogens control the pulsatile release of GnRH and can influence stress signaling (Acevedo-Rodriguez et al., 2018) and enhance HPA function (Handa et al., 1994). HSD17B11 is also related to CEBPB (CCAAT/enhancer-binding protein beta), a transcription factor regulating the expression of genes involved in immune and inflammatory responses (Chinery et al., 1997; Pless et al., 2008). The splicing of this gene, and others in our list, suggest a sex-specific response to stress mediated via alternative splicing.

As in our male-female splicing analysis, we identified few genes consistently alternatively spliced; however, the genes that were consistently spliced in both sexes were quite interesting. Rho family GTPases function as molecular switches to regulate a variety of cellular processes including cytoskeletal growth, gene transcription, and cytokinesis (Moon & Zheng, 2003; Miller & Bement, 2009). Both males and females displayed alternative splicing of the pituitary gene Rho GTPase Activating Protein 32 (ARHGAP32). Through response to GnRH pulsatile expression, Rho GTPases regulates cell motility and cytoskeletal rearrangements(Naor, Benard & Seger, 2000; Godoy, Nishimura & Webster, 2011). The shift from a control to stressed state likely places different constraints on cellular motility, requiring expedited movement of signaling molecules and energy transport throughout an organism; consistent splicing of ARHGAP32 may be one means by which cellular movement is adjusted during the stress response.

The hypothalamus of both males and females exhibit alternative splicing of the Retinitis Pigmentosa GTPase Regulator Interacting Protein 1 Like (RPGRIP1L) gene. RPGRIP1L is an adapter protein, localized at the ciliary transition zone, that interacts with G-protein coupled thromboxane A2 receptor (TBXA2R) to negatively regulate signaling at the ciliary base (Tokue, Sasaki & Nakahata, 2009; Gerhardt et al., 2015). This ciliary protein governs both autophagy and proteasome activity (Gerhardt et al., 2015; Struchtrup et al., 2018). As both sexes respond to stress by alternatively splicing the RPGRIP1L gene, the alteration of this locus may enable organisms to respond more effectively to a stressful environment through signal transduction and processing/disposal of damaged cells. Additional observation of the functioning of these gene products in control and stress states will provide further understanding as to the role these loci play in stress response.

### Control vs. Stress Splicing Comparison: Gene Ontology

In addition to characterizing splicing on a gene-by-gene basis, we assessed the gene lists, as a whole, to determine if a particular type of gene or genes related to a specific biological process were more readily spliced in the stress response. Our ontology analysis of the stress vs. control splicing events revealed elevated levels of splicing in genes with the following GO terms: small molecule binding, carb derivative binding, and drug binding (Figure 5). These terms were more abundant than genomic predictions in the male hypothalamus, female pituitary, and male gonad, while these same terms were less abundant than expected in the female hypothalamus and female gonad. Of the male tissues, the hypothalamus and testes appear more active in this area. There also were more spliced genes related to drug binding than expected.

**Figure 5.**
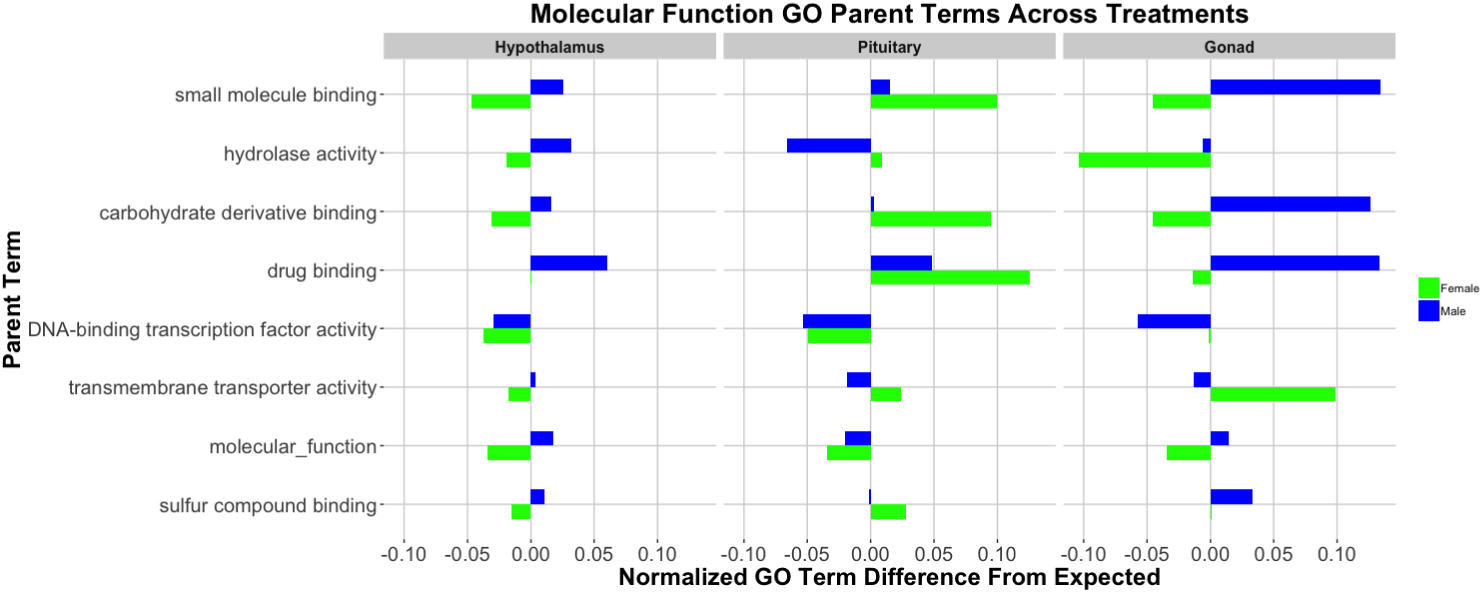
Molecular Function GO Analysis, Stress vs. Control (normalized counts of observed-expected). Binding genes were more frequently spliced in the male hypothalamus and the female pituitary. Counts for all terms in this figure were significantly different from the expected value in at least one of the tissues. Parent ontology terms along the y axis are in descending order from most frequent in the genome to least frequent. The three “panels” depict counts from the hypothalamus, pituitary, and gonads (respectively, from left to right). Abundances are observed counts – expected (based upon genomic predictions)/ total events within that tissue. We did not include any terms that were attributed to less than 2% of the genome.

### Control vs. Stress Splicing Comparison: Exon size, location, and motifs

Analysis of distributions of alternatively spliced exon sizes reveal that all tissues, regardless of sex, exhibit distributions of control-stress splicing events smaller than the genomic distribution (Wilcoxon, all p <2.80e-3)(Figure 3). The pituitary was the only tissue in which male and female distributions differed, with the male pituitary exons being longer than female (Wilcoxon, p = 0.022). Similar to the male-female splicing analysis of splicing in protein motifs, we found no significant difference in the distribution of events occurring in any one motif (Figure S6).

In analyzing locations of spliced exons, we again identified more events in coding regions than either 5’ or 3’ UTR, for all tissues (ChiSq p: hyp=4.41e-40, pit=9.60e-233, gon=5.13e-67)(Figure 6). Similar to our male-female analysis, we also identified a significantly lower abundance of splicing events in the 3’UTR regions of all tissues in the control-stress analysis (ChiSq p: hyp=0.042, pit=2.87e-12, gon=8.77e-4). There were fewer events in the 5’UTR of the pituitary (ChiSq p=2.04e-4), and more events occurred in the pituitary CDS than expected (ChiSq p=3.50e-4).

**Figure 6.**
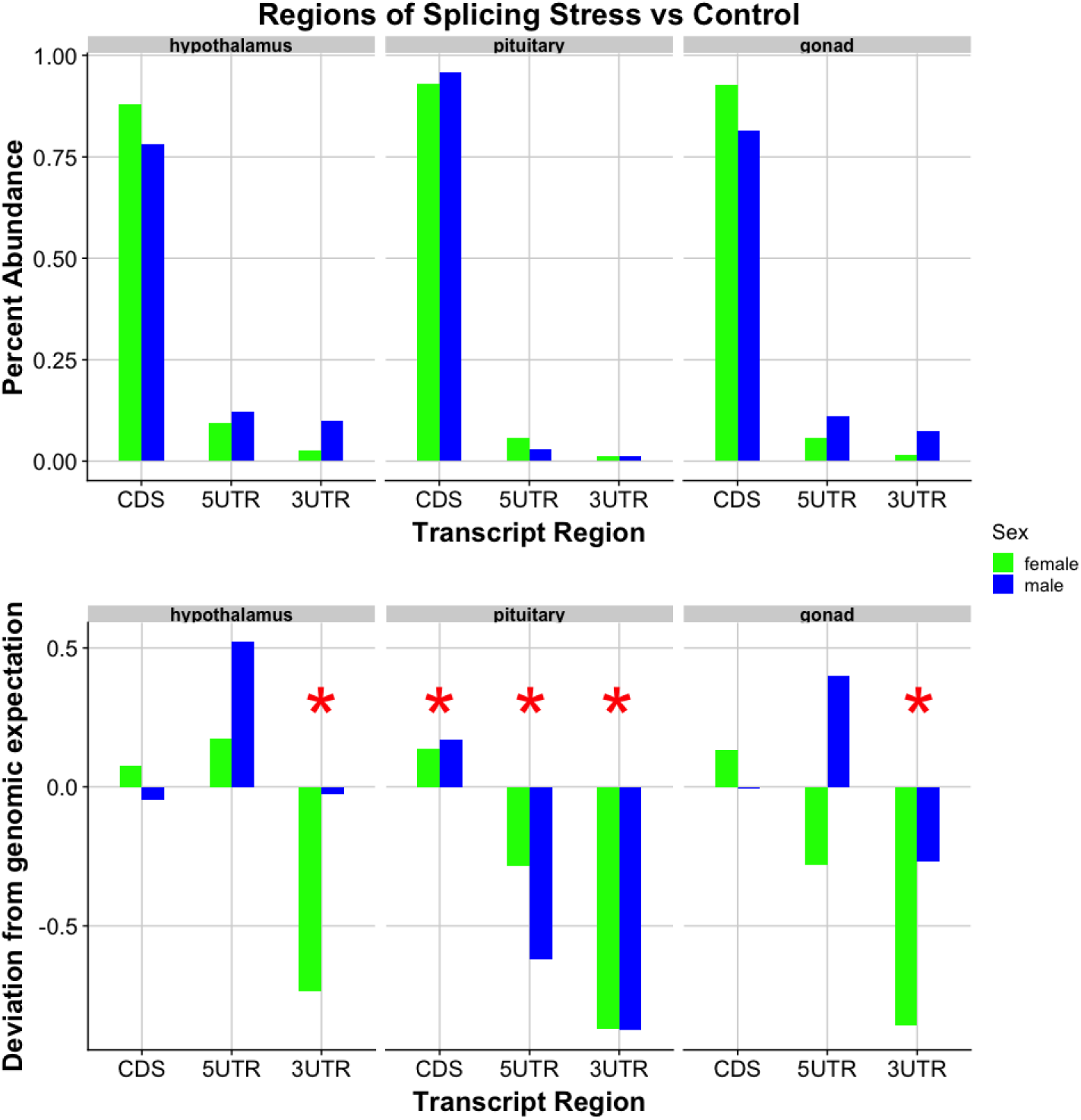
Transcriptional regions of control-stress splicing events. There were significantly fewer splicing events in the 3’UTR, agreeing with our previous findings, indicating that splicing machinery may selectively avoid this transcript region. The upper half represents percentage of total splicing events (for each tissue) in each transcript region. There were significantly more CDS events than 5’UTR or 3’UTR (ChiSq p: hyp=9.77e-23, pit=9.60e-233, gon=5.13e-67). The lower portion depicts how these counts differ from the expected (i.e. a value of 0.5 here indicates a 50% deviation from genomic expectation). Blue represents male, while green is female. Red stars represent significance between observed and expected proportions of splice location (ChiSq p: hyp-3’UTR=0.042, pit-CDS=3.50e-4, pit-5’UTR=2.04e-4, pit-3’UTR=2.87e-12, gon-3’UTR=8.77e-4).

### Treatment-specific and Sex-specific Isoforms

Because we know that treatment and sex affect gene expression documented (Calisi et al., 2018), we assessed each tissue for treatment-specific, and sex-specific isoforms of a gene. We did not identify any treatment-, nor sex-specific isoforms. This result suggests that even though gene expression may shift, and splicing events can be specific to an external stimulus, specific isoforms are not exclusive to one sex or treatment. A potential shortcoming of this particular analysis is the heavy reliance upon annotation. As the C. livia genome continues to improve with the progression of technology and sequence analysis tools we encourage others to reassess this work.

### Bimodal distribution of PSI

We assessed the distributions of PSI values for each splicing event type across all analysis comparisons to determine if there existed a differential in the frequency that a splicing event was present within a given state. The measure of “percent spliced in” reflects the frequency that one splicing isoform was seen over another in an individual. Ultimately, we find that PSI values do not show dramatic deviation for splicing events between treatments, nor between sexes (Figure S7). The only two instances that there appears to be a binary shift is in the case of alternate acceptors (AA), specifically in the hypothalamus of the male vs female analysis, and in the hypothalamus of the control vs stress analysis. This suggests that sex is not a driving factor for PSI values of stress-response alternative splice sites.

We would like to note that the algorithm used to identify significant splice events implicitly biases our results toward a bimodal distribution as any event with a deltaPSI between -0.1 and 0.1 was discounted. To determine how dramatically this shifted our distributions, we temporarily called all splicing events with p ¡= 0.05 significant. The additional 9 events this included for the male-female comparison were all found in the pituitary (3 control, 6 stress). Sixty-one events in the control-stress comparison had deltaPSI values between -0.1 and 0.1, 58 of which were also in the pituitary (53 male, 5 female) and 3 in the female gonad. Most significant splicing events have abs(deltaPSI) values greater than 0.1 in the male-female comparison. This is perhaps not particularly surprising as splicing differences are likely more biologically effective when included or excluded at higher PSI levels due to a more dramatic shift in protein presence for a given isoform. The 53 events in the male pituitary with low abs(deltaPSI) may suggest the male pituitary responds to stress not only by splicing genic regions at higher PSI values, but also by numerous, low-level splicing events. Further analyses using additional splicing tools and datasets are needed to validate this hypothesis.

## Discussion

To survive an unanticipated environmental perturbation, or stressor, the body activates multiple physiological systems to mobilize needed resources, with the intent of returning the body back to a homeostatic level (Michael Romero & Wingfield, 2015). The stress response is considered relatively conserved across vertebrates (Clarke et al., 1994; Tsigos & Chrousos, 2002; Leonard, 2005; Gutierrez-Triana et al., 2014; Taniuchi et al., 2016) and includes the limbic system, the central nervous system, the stress or hypothalamic-pituitary-adrenal (HPA) axis, and the thyroid or hypothalamic-pituitary-thyroid axis (HPT), all of which can directly or indirectly influence the HPG axis (Tsigos & Chrousos, 2002; Helmreich et al., 2005; Herman et al., 2005; Romero-Ramírez, Nieto-Sampedro & Barreda-Manso, 2017). Selection should favor an optimized stress response in which an organism’s physiological and behavioral responsiveness to perturbation promotes its survival and, ultimately, reproduction. However, we know little of how stress can affect the reproductive system at the genomic level of alternative splicing, and even less so about the similarities and differences experienced by males and females. Using the rock dove model in a highly-replicated RNA-Seq study, we tested how an external environmental stressor could affect the alternative splicing response of the HPG axis in both males and females. We discovered a significant sex difference in the splicing profile of hypothalamic and pituitary tissues. When subjects underwent a restraint stress treatment, we observed a decrease in the total number of significant splicing events between sexes as compared to controls (Figure 1). Two potential explanations for this are (1) the “convergent landscape hypothesis”: when males and females are stressed, they converge on a single splicing landscape, or (2) the “resource redirection hypothesis”: the stress response shuts down non-essential biological processes, minimizing background splicing “noise” that differentiates unstressed males and females, to redirect resources to bodily processes critical for survival.

### Convergent Landscape Hypothesis

The decrease in alternative splicing events experienced by stressed males and females as compared to their unstressed controls may suggest that both sexes are converging on an optimal splicing landscape. Previous reports show differential gene expression (DGE) between sexes in both an unstressed control state (Shapiro et al., 2013; Gillespie et al., 2013; MacManes et al., 2017), and in response to a stressor (Shapiro et al., 2013; Gillespie et al., 2013; MacManes et al., 2017; Calisi et al., 2018). Given these previous findings, coupled with ours presented here, we suggest that future study of the HPG stress response should endeavor to further characterize sex-specific and sex-convergent responses at all levels - genomic, proteomic, and methylomic. If other response mechanisms between males and females showed a similar pattern where the sexes exhibit fewer differences in a stress state, it may suggest that there was an optimal stress response phenotype being maintained at numerous levels, in addition to genomic ones currently understood. If this pattern of convergence is not seen at other physiological levels, it is possible that the decrease in splicing differences between males and females in the restraint stress treatment may reflect a stress response unique to the splicing level that does not exist in the proteomic or methylomic levels or additional variables unassessed in this study (e.g., stage of reproductive cycle).

### Resource Redirection Hypothesis

A second hypothesis to explain the presence of decreased alternative splicing differences between stressed males and females relative to controls is that the stress response redirects resources from less essential biological processes, like reproduction, to those more necessary for survival. Biological sex manifests in various distinct physiological phenotypes, particularly concerning those associated with reproductive processes (Ngun et al., 2011; Arnold & Itoh, 2011; Bundy, Vied & Nowakowski, 2017; Lee, 2018)However, an organism’s response to a stressor, no matter the sex, can trigger the mobilization of internal bodily resources to promote its survival (Wendelaar Bonga, 1997; Goldstein, 2010; Nicolaides et al., 2015; Smulders, 2017). Thus, an active sex-specific physiologically-driven alternative splicing landscape under stress may become reconciled in the HPG axis to shunt focus to other biological processes to support survival. This reduction in sexually dimorphic splicing background noise would result in fewer splicing differences between the sexes, as we observed. The remaining splicing differences may lie in loci generally implicated in the stress response, though future studies are needed to say this definitively. While our attempts to map spliced genes to Kegg pathways did not yield a discernable pattern of splicing of a particular pathway, we did find that several of the alternatively spliced genes may be involved in DNA-damage response and immune system processes, both of which are intimately linked with the stress response (Hara et al., 2011; Dhabhar, 2014). Additional study of the splicing landscape of males and females in various conditions, environments, age groups, and reproductive stages will shed further light on splicing variation between the sexes, and if our results seen are indeed a reflection of decreased activity of peripheral biological processes during stress.

### Genomic Events by Type

As compared to the hypothalamus and gonads, the pituitary in both sexes exhibited the most active splicing profile under stress as compared to controls. This pattern was consistent with our previous report of increased differential gene expression (DGE) in the pituitary following restraint stress (Calisi et al., 2018), suggesting this gland may be a targeted site for mediating a stress response in the HPG axis through both gene expression and alternative splicing mechanisms. Regarding overall splicing activity, males exhibited greater responsiveness to stress than females in the hypothalamus and pituitary, with a significant increase in response of the male hypothalamus compared to that of the female. Contrarily, examination of the same samples taken from the same birds showed that females exhibited greater DGE responsiveness to stress than males in all three tissues (Calisi et al., 2018). Thus, while examination at a gene transcription level may suggest the female reproductive axis as more responsive to the restraint stress treatment, examination at the alternative splicing level suggests males to be the more responsive of the sexes. The next steps would be to examine the protein products produced, but for now suffice it to say caution should be taken when determining which sex is more “responsive” to any stimuli, as this can vary by biological level of examination.

Further investigation of the types of splicing events that occurred following restraint stress revealed that core exon events as compared to all other event types were vastly more abundant across all tissues, regardless of sex or treatment, a phenomenon supported by findings in humans and mice (Koralewski & Krutovsky, 2011). Regarding the effects of stress treatment, more core exon events were present in the female gonads as compared to the male gonads. The higher abundance of core exon events could indicate that exon inclusion/exclusion is a more efficient approach to modulate a resulting protein, thereby triggering a more specific phenotypic response to a change in environment (Chen & Manley, 2009; Rodrigues, Grosso & Moita, 2013; Wang et al., 2015; Yabas, Elliott & Hoyne, 2015). This may then signify that the stress response of the female gonad is driven more heavily by core exon alterations to a transcript, while transcripts in the male gonad are more readily modified with all types of splicing events.

The only other splicing event type that differed significantly between the stress response of males and females was the retention of introns, with more retained intron events occurring in the male hypothalamus. Retained intron events are a means by which gene expression can be modified, often by downregulation (Ge & Porse, 2014). However, these events may also alter cis and trans elements that would otherwise modify transcript stability or translational efficacy (Jacob & Smith, 2017). Perhaps through differential utilization of the retained intron splicing event, the male hypothalamus could further modulate gene expression, targeting a more optimal stress response.

### Gene Ontology

In an attempt to gain a better understanding of potential products being generated by splicing activity, we examined our results using Gene Ontology (GO). Our inquiry of genes undergoing sex-biased alternative splicing revealed many GO terms related to “binding.” The function of these genes is linked with how hormones bind to their receptors, and by extension, how endocrine messages are received. With the increase in abundance of spliced hypothalamic binding genes and a decrease of “binding” terms in the pituitary spliced genes, it appears that males and females may differ in how cells in the hypothalamus bind various protein products. While the function of genes in the male and female pituitary related to “binding” exhibit fewer alternative splicing events than expected by genomic prediction, the sexes showed greater difference in alternative splicing of genes associated with “cell/neuronal structure”. This may suggest that similar functions can arise from sex-specific structures.

Alternative splicing exhibited in stressed versus control treatment subject HPG tissues also reveal sexual-dimorphic functionality. Once again, many of the alternatively spliced gene products within each sex in response to stress are related to “binding”. Specifically, we observed elevated splicing of these genes in the male hypothalamus, female pituitary, and male gonad. This pattern seems to suggest that the female pituitary is responding to restraint stress by altering how molecules/drugs are bound, while the male hypothalamus is more active in responding to stress via splicing binding genes. Thus, sexual differences in splicing landscapes in our samples may signify sex-biased hypothalamic function in response to stress, but similar pituitary functions between the sexes.

### Exon size, location, and motifs

The exon size distribution of alternative splicing events was significantly smaller than that predicted by the genomic distribution, agreeing with previous findings that the likelihood of a core exon being spliced may be inversely related to its size (Song et al., 2009). This inverse relationship of exon size and frequency of splicing may be a reflection of the highly-specific and precise adjustments required of splicing machinery (Black, 2000; Birzele, Csaba & Zimmer, 2008; Mittendorf et al., 2012). Inclusion or excision of a larger genetic sequence could prove more biologically challenging due to the physical distance between the two splice sites, and variable inclusion of a larger exon may result in a more dramatic alteration to the protein product, negating the effect of fine-tuning a response. Though the alternative splicing detected in both sexes exhibited smaller than expected exon sizes, size distributions in the female pituitary were smaller than those of the male pituitary. This may suggest that splicing in the female pituitary is targeted towards minute alteration, while genes in the male pituitary are modulated by movement/removal of larger sequence segments. Additional study of splicing in pituitaries of both sexes will be required to clarify the biological implications of this sex-bias.

In all comparisons, we found that the vast majority of splicing occurs in the protein coding region, or coding sequence (CDS) of a gene. When we examine the frequency of splicing location in relation to the genomic expectation (assuming an even distribution of splicing across all genic regions), we see that variation from expectation actually lies in the untranslated regions. Analyses comparing males and females, within each treatment group, revealed increased splicing of 5’-UTR genes in both the hypothalamus and pituitary, and a >6% decrease in splicing of the 3’-UTR genes in the pituitary. Splicing differences identified within the HPG axis of each sex in stressed birds as compared to controls displayed lower 3’UTR events in all tissues, and lower 5’UTR events in the pituitary. The increased abundance of 5’UTR alternative splicing events in males as compared to female subjects suggests that differences in the 5’UTR region of transcripts exist between the sexes (regardless of control/stress state). Previous research has found that the 5’UTR can mediate translation of proteins, regulate expression, and even regulate some splicing events (Kramer et al., 2013; Álvarez et al., 2016; Ohta et al., 2018). Thus, differential splicing of the 5’UTR between males and females may be a sex-specific mechanism for modifying protein production. The one instance in which we see a decrease in splicing of 5’UTR events is the pituitary, where each sex exhibits fewer splicing events in response to stress. Males and females may differ in the 5’UTR, but both sexes seem to minimize splicing of this region in the pituitary in response to restraint stress. The decrease in splicing events in the 3’UTR could signify that splicing events in the HPG axis do not favor the 3’UTR in response to restraint stress. 3’ ends of transcripts may be protected from alternative splicing, as evidenced by this decrease observed in both sexes and treatments. Degradation of mRNA transcripts can increase due to environmental stress (Canadell et al., 2015), and it may be that splicing events in the 3’UTR would result in undesired degradation of transcripts, or perhaps no degradation at all.

We did not identify any differences in the frequency of protein motif splicing between the sexes or treatments. As motifs combinations and conformations heavily impact protein function, we anticipated identifying specific motifs wherein splicing more readily occurred. Our query did not reveal this, suggesting that splicing is not targeting protein motifs within the HPG axis in either sex or treatment. Instead, the genic region (CDS, 5’UTR, 3’UTR) and gene-type appear to be larger drivers of the location that splicing events occur.

## Summary

In both sexes and treatment groups, we identified the most alternative splicing activity in the pituitary as compared to the hypothalamus and gonads. Our previous research also revealed the pituitary to have an increased response to restraint stress at the level of gene expression as compared to the hypothalamus or gonads (Calisi et al., 2018), suggesting a heightened stress response in this part of the HPG axis. Of genomic splicing events identified, core exon events are the most abundant form of splicing in all tissues/sexes/treatments, and splicing more frequently occurs in CDS regions. We found less splicing in the 3’UTR and more splicing in the 5’UTR than expected in both sex and treatment comparisons, suggesting that the 3’ end of transcripts is more biologically constrained than the 5’ end in our samples. Another splicing constraint evidenced by our findings, is that of an inverse relationship of exon size and splicing frequency. The overall reduction in splicing differences between the sexes when animals experienced a restraint stress treatment may point to a conserved splicing response to stress within the HPG axis. However, despite this reduction in sex-splicing differences, the male hypothalamus and female ovary experienced increased splicing activity in the face of stress as compared to the female hypothalamus and male testes, respectively. Sex differences in alternative splicing within the HPG axis, as well as in previously reported gene expression (Calisi et al., 2018) support sex-specific mechanisms for the stress response of the reproductive axis. While females experience increased differential gene expression in their HPG axis as compared to males in response to restraint stress (Calisi et al., 2018), we found that males experience increased alternative splicing. Our examination of the vertebrate stress response at multiple levels of biological organization offers a more complete picture of its mechanistic underpinnings between the sexes at an unprecedented proximate level. These data inspire further integrative levels of investigation to inform and potentially revolutionize evolutionary theory and the study of stress-induced reproductive dysfunction.

## Competing interests

The authors declare that they have no competing interests.

## Availability of data and materials

Read data corresponding to the control birds are available using the European Nucleotide Archive project ID PRJEB16136; read data corresponding to the stressed birds are available at PRJEB21082. All code for analyses in this manuscript can be found at https://github.com/AndrewLangvt/Scripts/tree/master/splicing_analysis/.

## Funding

This work was funded by NSF IOS 1455960 (to RMC and MM).

## Author’s contributions

RMC and MM designed the study. SA conducted avian manipulation and tissue sampling. AL conducted RNA extraction, cDNA library preparation. AL conducted computational analyses with input from RH. AL wrote the manuscript with input from all.

## Acknowledgements

We thank the many undergraduate and graduate students of the MacManes and Calisi Lab for their assistance on this project, including their involvement in avian husbandry, conducting the experiment, tissue collection, RNA extraction, and cDNA library preparation.

## Supplementary Files

Table S1. List of Spliced Genes.

https://docs.google.com/spreadsheets/d/1FqrZGmNXVwxxQN-nDb4ML25Vo8b-lsud0OW5JZjMAus/edit#gid=0 This list contains the Entrez ID for all genes in which an alternative splicing event was found during this study. Column headers refer to the comparison (M= male, F= female, H= hypothalamus, P= pituitary, G= gonad, C= control, S= stress)

**Figure S1.**
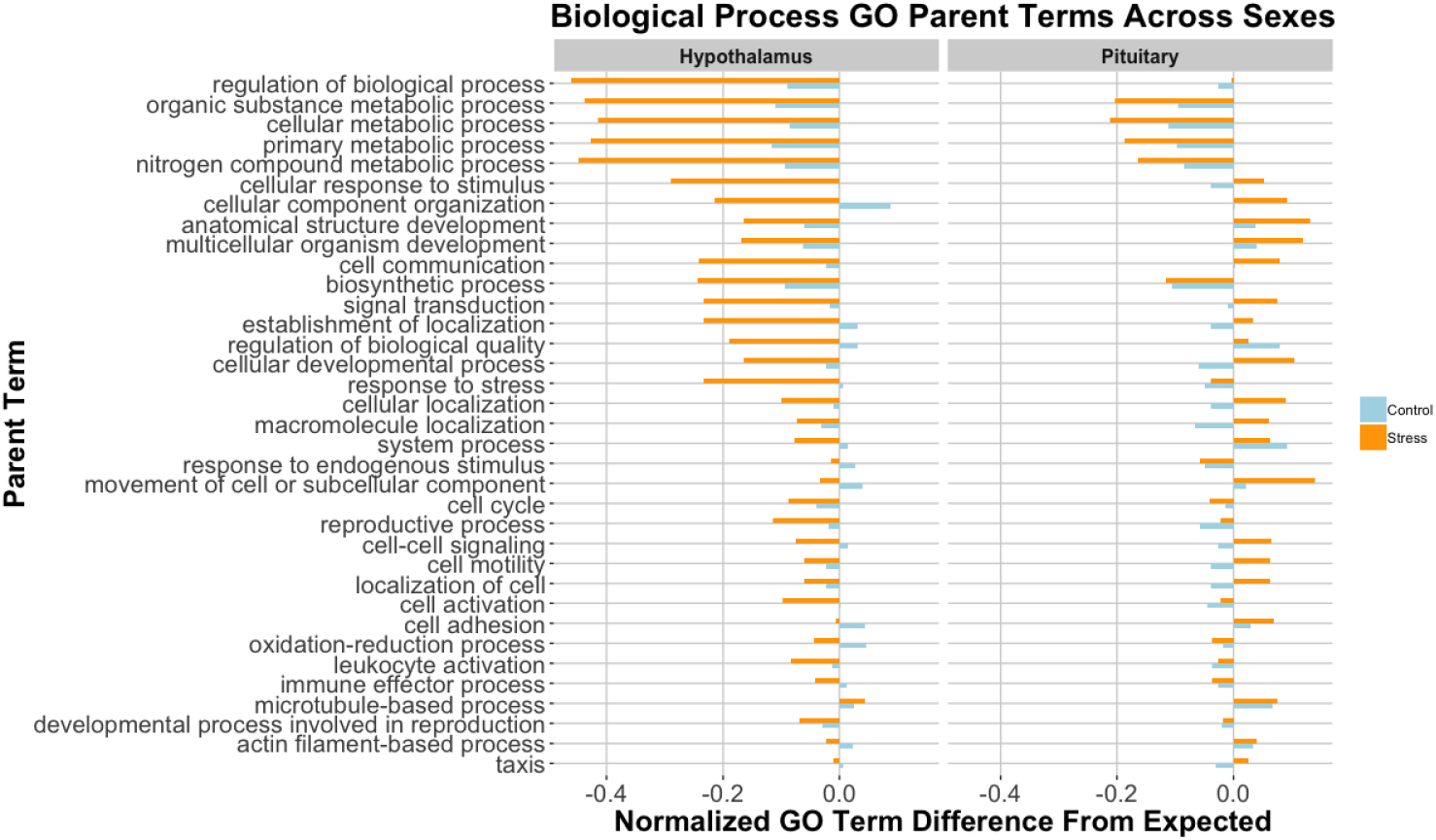
Male vs. Female Biological Process GO Analysis

**Figure S2.**
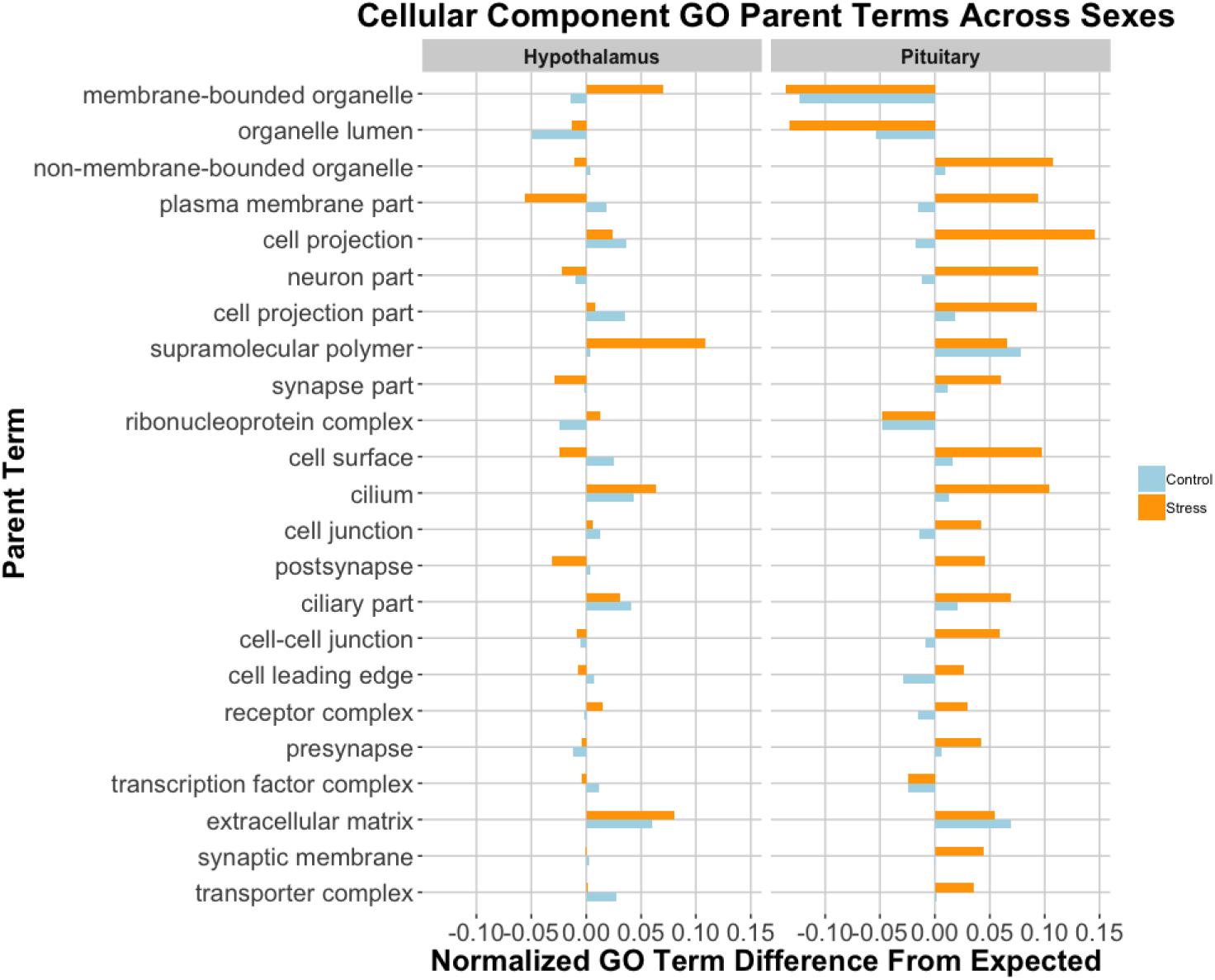
Male vs. Female Cellular Component GO Analysis

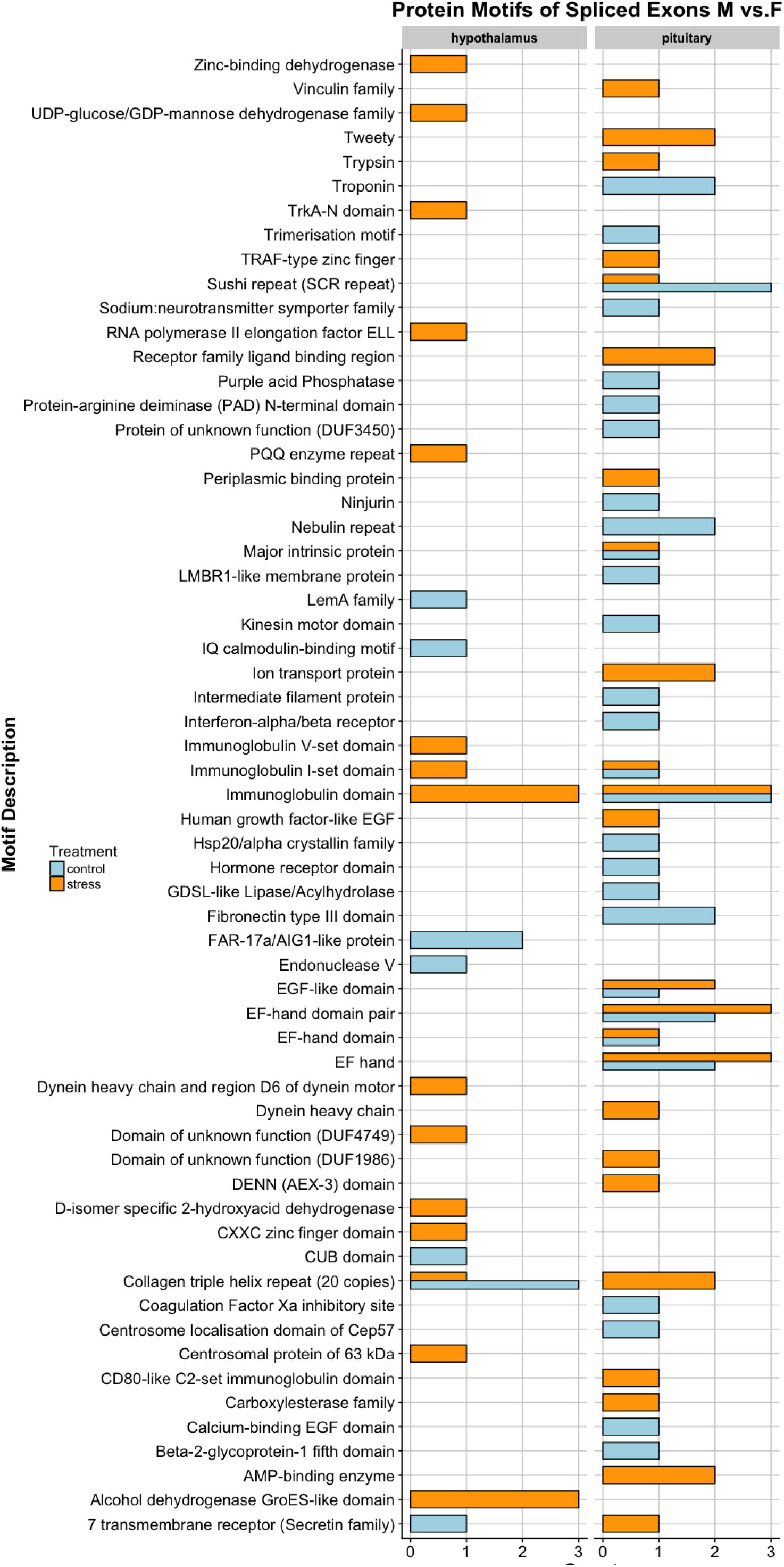

**Figure S4.**
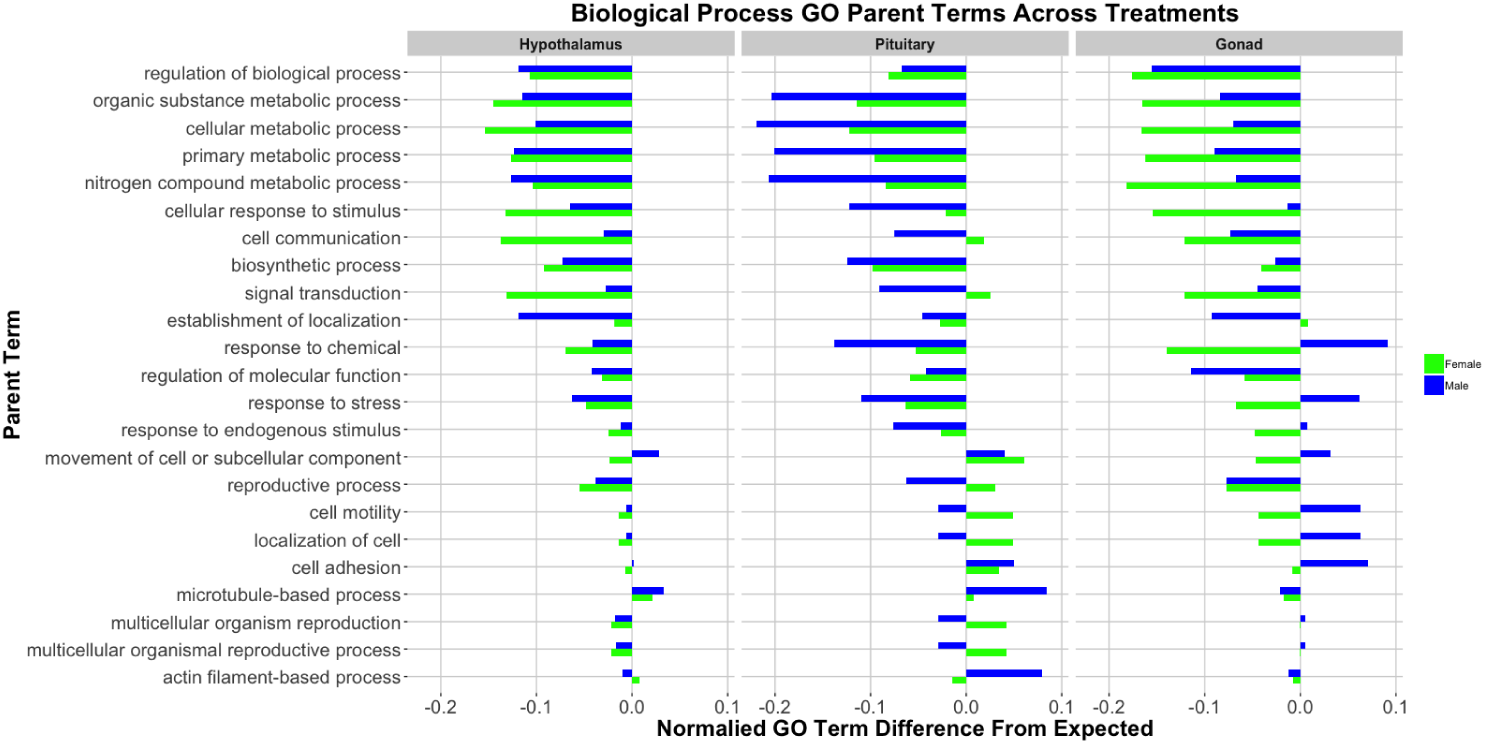
Control vs. Stress Biological Process GO Analysis

**Figure S5.**
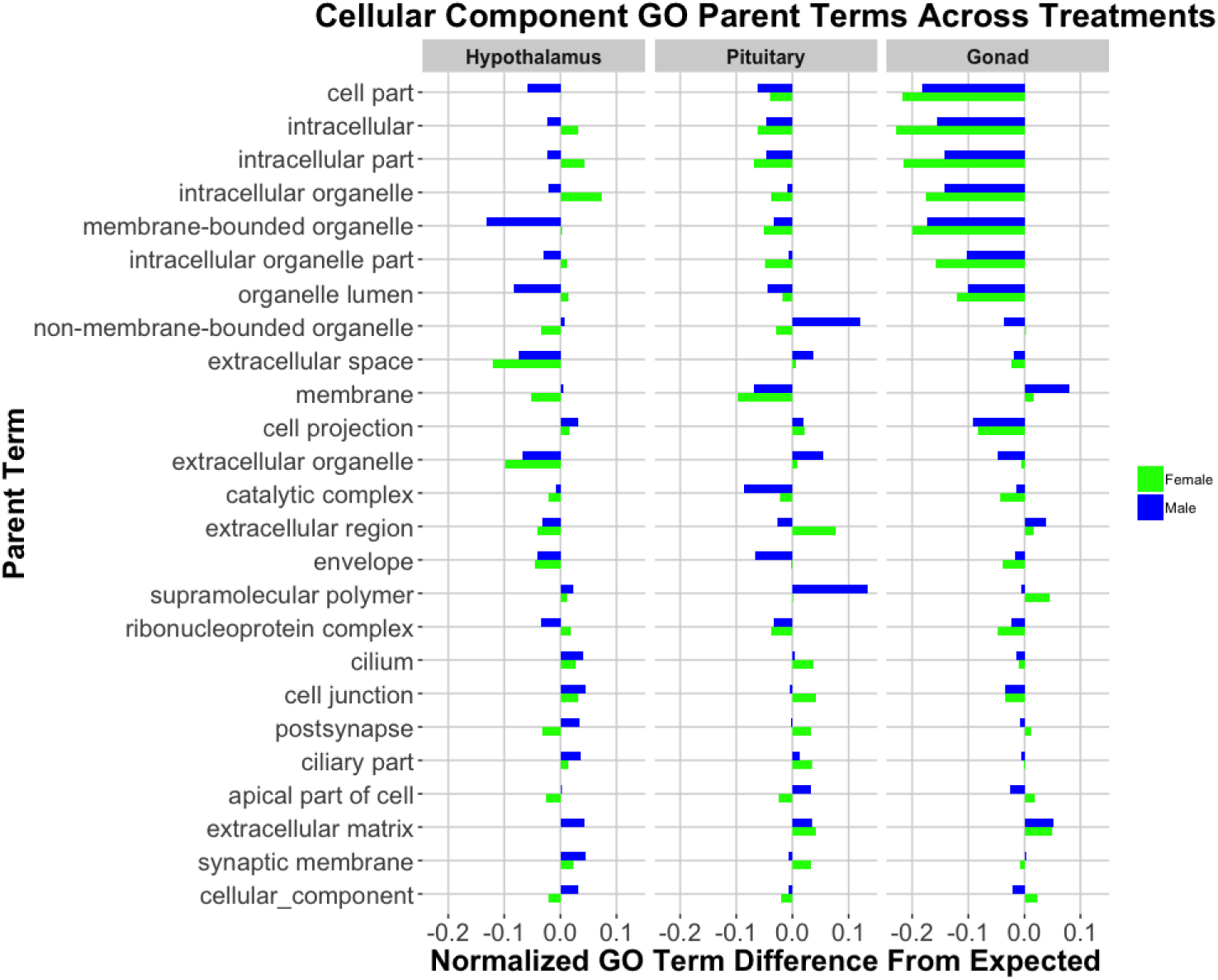
Control vs. Stress Cellular Component GO Analysis

**Figure S6.**
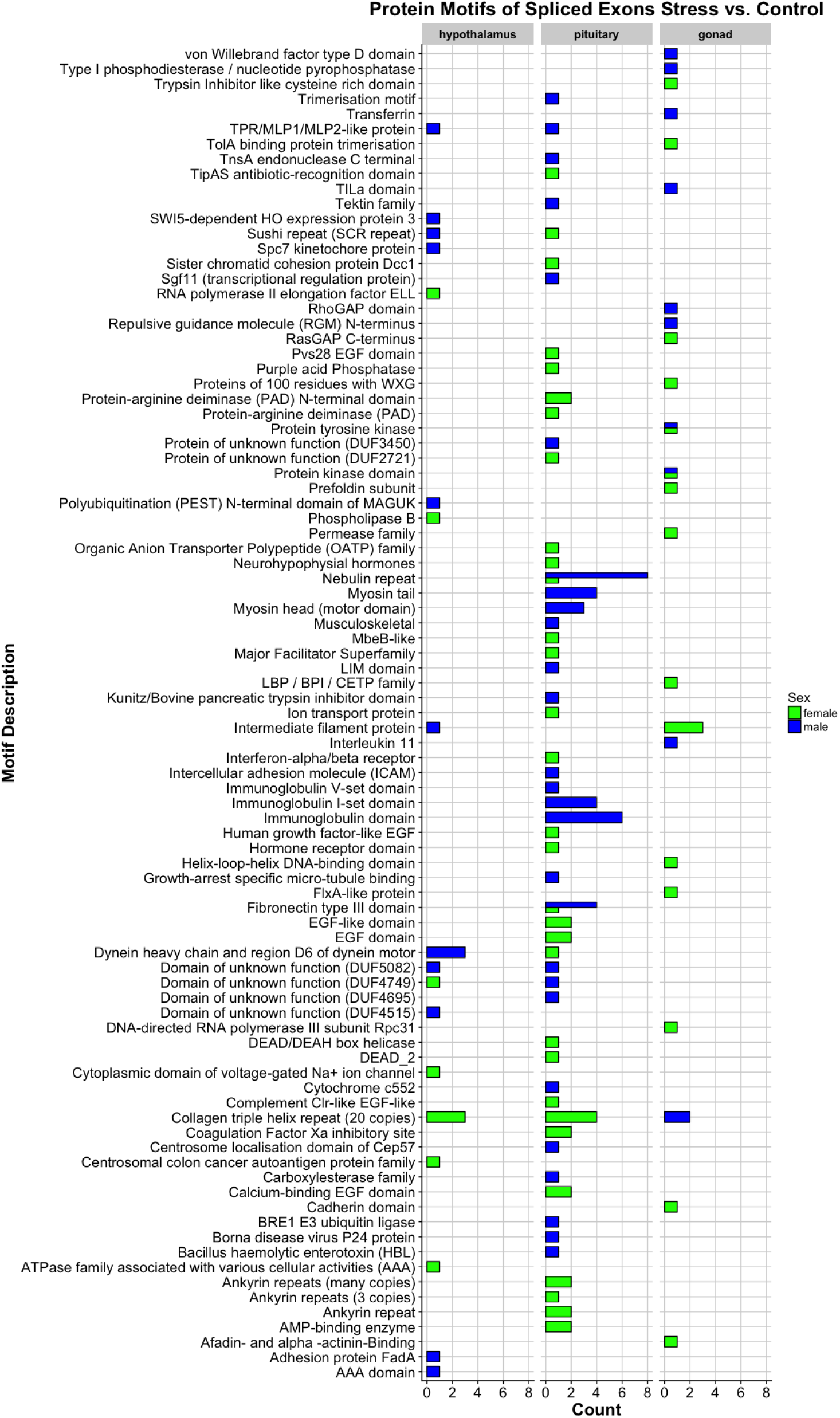
Control vs. Stress Exon Motifs

**Figure S7.**
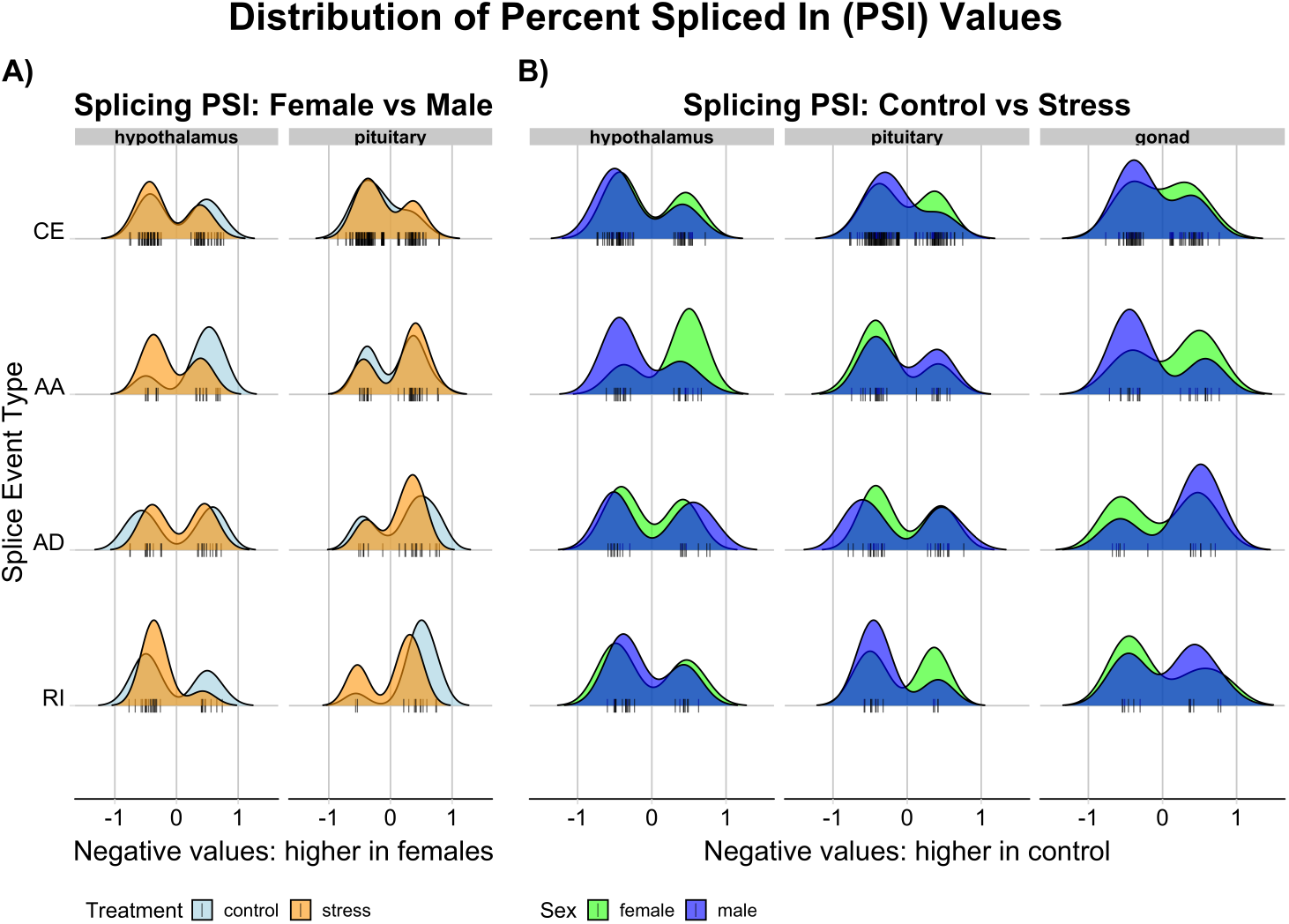
Distribution of Percent Spliced In (PSI) Values for the a) control vs stress analysis and also the b) male vs female analysis. The figure is divided into vertical panels, each representing the tissues labeled a the top The right two panels show the distribution of Percent Spliced In (PSI) values, with “—” symbols at the base of each ridge plot each corresponding to the PSI value of a single event. The heights of the ridge plots should not be compared from one event to another, but within an event between treatments. Splicing event types are displayed along the y-axis (CE: core exon, AA: alternate acceptor, AD: alternate donor, RI: Retained intron). The coloring scheme is the same as previous figures (light blue: control, orange: restraint stress, green: female, blue: male).

## Notes

#### Summary of Updates

formatting updated to enhance readability

